# OEFinder: A user interface to identify and visualize ordering effects in single-cell RNA-seq data

**DOI:** 10.1101/025437

**Authors:** Ning Leng, Jeea Choi, Li-Fang Chu, James A. Thomson, Christina Kendziorski, Ron Stewart

## Abstract

A recent paper identified an artifact in multiple single-cell RNA-seq (scRNA-seq) data sets generated by the Fluidigm C1 platform. Specifically, Leng* *et al*. showed significantly increased gene expression in cells captured from sites with small or large plate output IDs. We refer to this artifact as an ordering effect (OE). Including OE genes in downstream analyses could lead to biased results. To address this problem, we developed a statistical method and software called OEFinder to identify a sorted list of OE genes. OEFinder is available as an R package along with user-friendly graphical interface implementations that allows users to check for potential artifacts in scRNA-seq data generated by the Fluidigm C1 platform.

**Availability and Implementation:** OEFinder is freely available at https://github.com/lengning/OEFinder

## Introduction

Single cell RNA-seq (scRNA-seq) has led to important findings in many fields and is becoming increasingly popular in studies of transcriptome-wide expression (Deng* *et al*. 2014; Shalek* *et al*. 2014; Trapnell *et al*. 2014; Treutlein* *et al*. 2014; Leng* *et al*. 2015). To facilitate scRNA-seq, the majority of studies ultilize the Fluidigm C1 platform for cell capture, reverse transcription, and cDNA amplification, as this platform allows for rapid and reliable isolation and processing of individual cells. In spite of the advantages, Leng* *et al*. (2015) identified an artifact in multiple data sets generated by C1 and confirmed that the artifact was present in the cDNA processed by the C1 machine. Specifically, in these datasets, there are genes showing significantly higher expression in cells captured in capture sites with small or large plate output IDs. We refer to this artifact as an ordering effect (OE), which has been shown to be independent of organism and laboratory (Leng* *et al*. 2015). As detailed in Leng* *et al*. (2015), accurate identification of OE genes is important to ensure unbiased downstream analyses.

Leng* *et al*. (2015) used an ANOVA-based approach to detect OE genes. The ANOVA-based approach performs well in many cases, but has reduced power when few cells are available. (Note that in empirical data, the cells can be missing by the random effects of capture failure - an empty capture site, or doublets - capturing more than one cells in one capture site). To improve power for identifying OE genes, we developed an approach, OEFinder, based on the orthogonal polynomial regression. Results show that OEFinder is less sensitive to sample size and outperforms the ANOVA-based approach when few cells are available.

OEFinder is implemented in R, a free and open source language, with a vignette that provides working examples. The graphical user interface (GUI) impelementations of OEFinder allow a user with little computing background to easily identify and characterize OE genes in scRNA-seq data (Figure 1a and Supplementary Figure 4).

**Figure 1.**
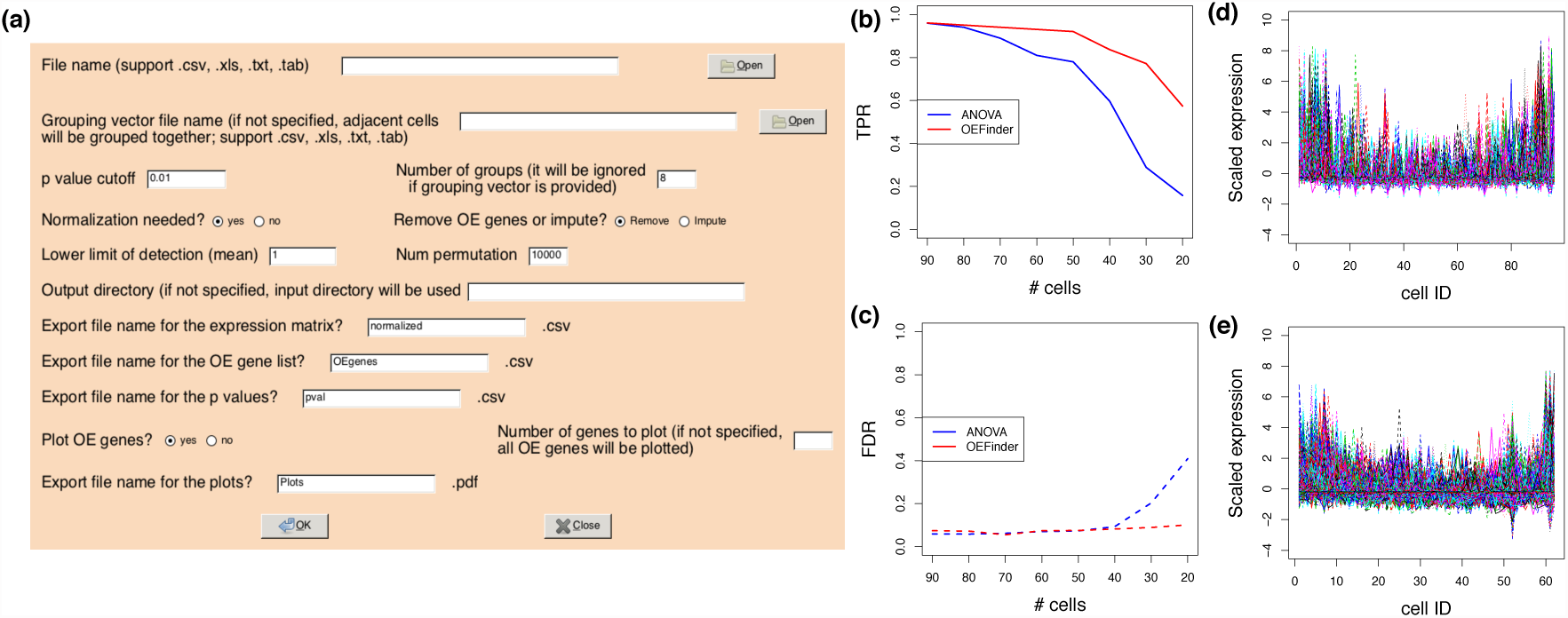
Panel (a) shows the OEFinder GUI for identifying OE genes (shown is implementation using R/RGtk2 package; an implementation using R/shiny package is also available, see Supplementary Figure 4). Panels (b) and (c) show the number of available cells. The y-axis shows true positive rate (TPR) and false discovery rate (FDR). Panel (d) and (e) show the OE genes identified in the first experiment of Trapnell *et al.* data and Leng* *et al.* data, respectively. The cells were ordered following the capture site ID. The y-axis shows scaled gene expression (z-score). Each line represents one OE gene.

## Analysis input and output

### INPUT

#### Expression estimates

OEFinder requires a genes-by-cells expression matrix. The expression matrix can be either normalized or unnormalized. If the input matrix is unnormalized, OEFinder applies the Median-by-Ratio normalization method introduced by Anders and Huber (2010) prior to OE detection.

#### Capture site group definitions

As detailed in Leng* *et al*. (2015), the capture sites are labeled as A01, …, A12, B01, …, B12, …, H01, …, H12. If the capture site IDs are provided, OEFinder groups cells from sites with the same starting letters. When the capture site information is not available, OEFinder groups cells based on their input order. By default, OEFinder groups cells into 8 even-sized groups. The number of groups may be changed by a user.

### METHOD

The normalized expression values of each gene are scaled to z-scores. For each gene, OEFinder applies an orthogonal polynomial regression on z-scores against group code. To infer whether gene *g* follows the OE trend, OEFinder calculates the p-value *p*_*g*,2_ of a one-tailed test that tests whether the coefficient of the quadratic term is positive. To account for the goodness of the spline fitting, OEFinder defines an aggregate statistics *S_g_* as -log(*p*_*g*,2_)-log(*p_model_*), in which *p_model_* denotes the F test p-value of the full model. OEFinder then generates 10000 simulated genes from permuted data to evaluate the significance of the observed aggregated statistics. By default, genes with permutation p-value less than 0.01 are identified as OE genes. The number of simulated genes and the p-value cutoff may be changed by a user (for further details, see Supplementary section 2).

### OUTPUT

#### List of OE genes

OEFinder outputs two .csv files - one contains a sorted list of OE genes and the other contains p-values for all genes.

#### Expression matrix for downstream analysis

OEFinder outputs a normalized expression matrix that can be directly input to downstream analyses. The user has the option to choose either removal of the OE genes, or imputation of the OE genes with adjusted values.

#### Visualization of OE genes

OEFinder generates a .pdf file contains expression plots of the top N OE genes, where N is user-specified. An example is shown in Supplementary Figure 1.

## Evaluations

### Simulation studies

We conducted 8 simulation studies to evaluate the performance of the OE detection algorithms. In each simulation, we generated 5000 expressed genes with 500 OE genes. Expression of OE genes was generated based on expression profiles of OE genes detected in empirical data (Details of the simulations may be found in Supplementary section 3). The 8 simulation studies evaluate cases with varying numbers of available cells (20-90 cells). Each simulation study contains 100 repeated simulations.

Figure 1b-c show the true positive rate (TPR) and false positive rate (FDR) comparing the ANOVA-based method introduced in Leng* *et al.* (2015) and OEFinder. Results indicate that when more than 60 cells are available, both methods have TPR greater than 90% while FDR is controlled below 10%. When fewer cells are available, OEFinder has a higher TPR than the ANOVA-based approach and the FDR is still well controlled. Additional simulation results may be found in Supplementary section 4.

## Case studies

We applied OEFinder on two publicly available data sets with capture site ID information. Trapnell *et al.* (2014) data and Leng* *et al.* (2015) data contain 4 and 3 experiments, respectively. Figure 1d-e show 187 and 451 OE genes identified in the first experiment of each data set. Genes detected by OEFinder show a clear OE pattern. Results of other experiments may be found in Supplementary section 5.

## Discussion

We developed an R package OEFinder which can robustly detect OE genes in scRNA-seq data generated by Fluidigm C1. OEFinder provides user-friendly graphical interface implementations that facilitate use by investigators with little computing backgrounds.

